# Characterization of the SKC mouse strain as a potential model for keratoconus

**DOI:** 10.64898/2026.04.27.721145

**Authors:** Rachel Hadvina, Jingwen Cai, Hongfang Yu, Amy Estes, Yutao Liu

**Author notes:** **Corresponding author:** Yutao Liu, Department of Ophthalmology, Mayo Clinic, Rochester, MN 55902.

## Abstract

**Background:** Keratoconus (KC) is a multifactorial disorder with unclear etiology, characterized by localized thinning and a cone-like protrusion of the cornea. The complex etiology of KC exacerbates the lack of an animal model. Previous studies by Tachibana et al. (2002) described an inbred mouse strain (SKC) with a spontaneous, androgen-dependent, cone-like corneal morphology. This study aimed to investigate the corneal phenotypes of SKC mice through an in-depth ophthalmic examination.

**Methods:** Mice (n=53) were examined via slit lamp biomicroscopy with fluorescein staining. Spectral-domain optical coherence tomography (SD-OCT) enabled central corneal thickness (CCT) measurement in selected mice (n=26 eyes), and OCT-based pachymetry mapping (n=16 eyes). In vivo corneal confocal microscopy was conducted on eyes to assess cellular morphology (n= 9 eyes). Eyes were collected for histology analysis (n=22).

**Results:** Lesions and epithelial breaks were present in ∼95% of eyes (n=101). Neovascularization, perforation, scarring, and hydrops were seen primarily in males. An opaque, unilateral cone-like morphology was exclusive to males (n=11). Male and female corneas showed no significant difference in CCT, though pachymetry mapping revealed regional thinning patterns in both sexes. Loosened epithelial tight junctions, stromal fibrosis, vascularization, and inflammation of variable severity were identified in both sexes.

**Conclusion:** This study identified previously unreported corneal phenotypes in SKC mice through ophthalmic examination. Unlike previous studies, gross and histological abnormalities were observed in female SKC mice. Our findings suggest a lower penetrance of the cone-like phenotype (∼20%) than previously reported (∼33%) and support that the conical phenotype in male mice may be secondary to keratitis.

## Introduction

Keratoconus (KC), the most common corneal ectasia disorder, is characterized by bilateral, asymmetric corneal degeneration with local, centralized thinning and protrusion of the cornea [1]. KC often manifests at puberty, affects both sexes, and occurs within different ethnic backgrounds with no well-established bias [1-3]. KC affects an estimated 0.2 to 4,790 per 100,000 people, with prevalence rates varying by population and more recent estimates suggesting even higher rates [2, 4-7]. Symptoms of KC are progressive with myopia and irregular astigmatism at the initial stages and worsening corneal steepening and corneal scarring at more advanced stages [8]. Patients experience vision impairments and reduced quality of life, which worsen as the disease progresses. Treatments for KC are limited. Early, conservative treatments aim to correct refractive errors and improve visual acuity [3]. However, these treatments do not slow or halt disease progression. The most modern treatment for KC, corneal collagen crosslinking (CXL), is used to crosslink collagen fibrils in the corneal stroma, strengthening the cornea and mitigating further progression of KC [9, 10]. However, CXL cannot be performed in patients with severe thinning, scarring, or corneal opacification [11]. Currently, surgical intervention, namely full corneal transplantation, is the most effective form of treatment for severe KC [2]. KC is widely understood to be a multifactorial disorder, with factors believed to impact KC pathogenesis, including genetic variants, environmental stressors, biomechanical stressors, hormonal changes, atopy, and other comorbidities [12-18]. However, the precise etiology of KC pathogenesis remains unclear.

The human cornea is composed of five main layers: the epithelium, acellular Bowman’s layer, collagenous stroma, acellular Descemet’s membrane, and endothelium. The corneal epithelium is the anterior-most layer and is composed of a non-keratinized stratified squamous cell layer [19]. The stroma, a layer of well-organized collagenous lamellae and keratocytes, is crucial for the transparency and structural integrity of the cornea [20, 21]. Stromal keratocytes contribute to maintenance of the extracellular matrix (ECM) environment and stromal homeostasis [22]. The epithelium accounts for around 10% of the total corneal thickness, whereas the stroma accounts for around 80% of corneal thickness [19, 21]. Histopathological changes associated with keratoconus include stromal thinning, Descemet’s membrane rupture, stromal or corneal scarring, and epithelial changes [2]. These histopathological changes are the driving factors of KC-associated corneal ectasia [23].

Due to the complex nature of KC, an animal model that replicates both underlying molecular mechanisms and the cone-like corneal phenotype has not been established [24, 25]. Prior studies have attempted to generate animal models by genetic modification and by applying chemicals or enzymes to the cornea [24, 25]. One set of studies by Tachibana et al. (2002) and Quantock et al. (2003) described a strain of inbred mice (SKC) with a spontaneous, androgen-dependent cone-like corneal morphology [26, 27]. The SKC mice exhibited a cone-like, steepening, and bulging of the eyes that varied in severity among mice and was transmitted between generations in an autosomal recessive manner with around 33% penetrance and a bias for males [26]. Cell growth, apoptosis, and c-fos expression, a KC-associated transcriptional regulator, were assessed in the original publication [26]. Electron microscopy recapitulated gross observation by demonstrating a drastic elevation in corneal surface curvature [26]. Hematoxylin & eosin (H&E) staining revealed abnormal central corneal thinning, curvature, and steepening in male and female SKC mice injected with testosterone [26]. Chromosomal linkage mapping of this corneal phenotype identified a locus on mouse chromosome 17 within the major histocompatibility complex (MHC) region, with a logarithm of the odds (LOD) score of 9.77 assigned to markers *D17Mit32* and *D17Mit34* [26]. Quantock et al. (2003) assessed the microstructures of the SKC mouse cornea and found the collagen fibrils to be larger and more widely spaced than the wildtype mouse cornea [27]. Despite these examinations, a full, in-depth ophthalmic examination of the SKC corneal phenotype has not been conducted, limiting the utility of the strain for disease modeling. This study sought to conduct an in-depth ophthalmic examination of SKC mice to characterize precise corneal phenotypes and assess their utility as an experimental model for keratoconus.

## Methods

### Animal husbandry

All experimental protocols were approved by the Institutional Animal Care and Use Committee (IACUC) at Augusta University (protocol number 2014-0697) and were conducted in accordance with the Association for Research in Vision and Ophthalmology (ARVO) Statement for the Use of Animals in Vision and Ophthalmic Research. SKC mice (5 females and 4 males) were acquired from the RIKEN BioResource Research Center (Koyadai, Tsukuba, Ibaraki, Japan) after cyrorecovery. Mice were group-housed in a temperature and humidity-controlled conventional housing facility under a 12h/12h light/dark cycle. A standard diet was provided ad libitum, with automatic water access, and health surveillance overseen by the Augusta University Division of Laboratory Animal Services, with veterinary oversight. Ophthalmic examination occurred between 2 and 7 months of age, with an average age at examination of ∼3.7 months.

### Mouse phenotyping

All mouse phenotyping, including slit lamp biomicroscopy, spectral-domain optical coherence tomography (SD-OCT), and in vivo cornea confocal microscopy, was conducted with the support of the Augusta University Medical College of Georgia Visual Function Assessment Core Facility (RRID: SCR_027277).

### Slit lamp biomicroscopy

Mice (n=53) were lightly anesthetized with isoflurane USP (NDC 11695-6777-2, Covetrus North America, Dublin, OH, USA), which was administered with a vaporizer through an induction chamber and nose cone. Mice were then examined and imaged with slit lamp biomicroscopy using the ZEISS SL800 slit lamp (ZEISS Group, Oberkochen, Germany) at either 25X or 40X magnification. BioGlo fluorescein ophthalmic strips (HUB Pharmaceuticals LLC, Scottsdale, AZ) were used to assess breaks in epithelial tight junctions under a cobalt blue filter. Neomycin and polymyxin B sulfates and bacitracin zinc ophthalmic ointment, USP (NDC 24208-780-55, Bausch & Lomb Americas Inc., Bridgewater, NJ, USA) was applied to the eyes after each examination to prevent ocular dryness and mitigate infection risk.

### Spectral domain-optical coherence tomography

Anterior segment SD-OCT was used to visualize mouse corneal curvature and thickness in selected mice (n=13). The Bioptigen Envisu-R2200 Spectral Domain Ophthalmic Imaging System (Leica Microsystems, Wetzlar, Germany) was used with a 12mm telecentric probe with the reference arm set to position 648. Mice aged 2 to 7 months were anesthetized based on body mass (10 μL/g) with 100 mg/ml ketamine (NDC 11695-0703-1, Covetrus North America, Dublin, OH, USA) and 100 mg/mL xylazine (NDC 11695-4024-1, Covetrus North America, Dublin, OH, USA). Systane eye drops (Alcon, Fort Worth, TX, USA) were applied to the surface of the eye to clean the surface of the cornea of any debris and to prevent corneal dryness during examination. Each eye underwent one rectangular and one radial scan. Following the examinations, neomycin and polymyxin B sulfates and bacitracin zinc ophthalmic ointment, USP (NDC 24208-780-55, Bausch & Lomb Incorporated, Tampa, FL) was applied to each eye to prevent corneal dryness and mitigate infection while the mice recovered from the anesthesia. Central corneal thickness (CCT) was measured in each cornea using the caliper function in the InVivoVue Diver 2.4 software (Leica Microsystems, Wetzlar, Germany). Differences in CCT between males and females were assessed with GraphPad Prism v. 10.5.0 using a two-tailed t-test at an alpha of p<0.05.

### In vivo cornea confocal microscopy (IV-CCM)

Cell-containing layers of the mouse cornea were examined using the HRT3-RCM in vivo cornea confocal microscope from Heidelberg Engineering (Heidelberg Engineering Inc, Franklin, MA). Mice were anesthetized prior to examination with ketamine and xylazine based on body mass as previously detailed. Systane eye drops (Alcon, Fort Worth, TX, USA) were first used to clean the surface of the eye of any debris. GenTeal eye gel (Alcon, Fort Worth, TX, USA) was then applied to both eyes to prevent dryness between examinations and to allow imaging. GenTeal (Alcon, Fort Worth, TX, USA) eye gel was also applied to the interior of a sterile TomoCap and the surface of the microscope lens (Heidelberg Engineering Inc, Franklin, MA). The mice were positioned so that the center of the eye was lined up with the center of the microscope lens. The microscope was then moved forward to make gentle, direct contact with the eye. The focal depth of the microscope view was adjusted to allow viewing of the epithelium, with a depth setting of 0 µm. High-resolution images of the epithelium, stroma, and endothelium were obtained using section scans of the region of interest at a single depth and volume scans, which capture 40 images of the region of interest with 2 µm spacing between images. Following the examinations, neomycin and polymyxin B sulfates and bacitracin zinc ophthalmic ointment, USP (NDC 24208-780-55, Bausch & Lomb Incorporated, Tampa, FL) was applied to each eye to prevent corneal dryness and mitigate infection while the mice recovered from the anesthesia. Epithelial and endothelial cell density (cells/mm^2^) was calculated using the semi-automatic cell counting function built into the microscope. Epithelial and endothelial cell density were compared between males and females with GraphPad Prism v.10.5.0 using a two-tailed t-test at an alpha of p<0.05.

### Pachymetry mapping

OCT-based corneal pachymetry mapping was conducted using the mouse corneal analysis program (MCAP) software v.1.0 [28]. Maps were generated from radial scans of each cornea. The epithelial and endothelial borders were defined by a semi-automated, gradient-based algorithm applied to boundaries manually drawn at the 50^th^ of 100 frames [28]. The remaining frames of the scan are automatically segmented by the software, and axial motion correction is applied [28]. To create 3D representations of each corneal surface, corrected data points are resampled to a uniform Cartesian distribution, and a Zernike polynomial expansion is then used [28]. Pachymetry maps were created by subtracting the z-axis of the epithelial and endothelial borders. The resulting maps are displayed as a heatmap to visualize regions of corneal thinning or thickening.

### Histology and immunofluorescence

Eyes (n=22) were collected at 3-6 months of age, fixed in Davidson’s fixative for 24 hours, and immediately transferred to 70% ethanol until further processing. Eyes were embedded in paraffin and sectioned with a thickness of 5μm on a sagittal plane through the optic nerve head to the center of the cornea. Three sections per eye were processed for H&E staining, and three sections from selected eyes (n=8) were processed for immunofluorescence (IF) staining at the Augusta University Medical College of Georgia Electron Microscopy and Histology Core Facility, RRID: SCR_026810. Corneal tissues selected for IF were double-stained with a combination of either zona occludens-1 (ZO-1) (1:100, Invitrogen #61-7300) and α-smooth muscle actin (α-SMA) (1:100, Invitrogen #14-9760-82) or claudin-2 (1:100, Invitrogen # 32-5600) and superoxide dismutase-2 (SOD2) (1:100, Invitrogen #PA5-30604). Additionally, cell nuclei were stained with 4’,6-diamidino-2-phenylindole (DAPI). H&E slides were imaged with brightfield microscopy at the AU Cell Imaging core using the Zeiss Axioscan 7 (ZEISS Group, Oberkochen, Germany) slide scanner set at 10X magnification and processed using the ZEISS ZEN Microscopy software v.3.10. IF slides were imaged via confocal microscopy in the Augusta University Medical College of Georgia Cell Imaging Core Facility, RRID:SCR_026799, using the Leica Stellaris 5 confocal microscope (Leica Microsystems, Wetzlar, Germany) at 20x and 63X magnification.

## Results

### Corneal morphology

Changes in the gross corneal morphology, including changes in shape, curvature, clarity, and other clinical pathological features, were observed with slit lamp biomicroscopy (Figure 1A). The most prominent change to the corneas of male and female SKC mice was epithelial lesions and defects, with ∼95% of the eyes (n=101) demonstrating epithelial abnormalities upon gross observation and fluorescein staining of the corneas (Figure 1A &B). A unilateral cone-like morphology accompanied by opacity was identified in 11 male mice (n=11 eyes). The severity of the protrusion varied among mice. Among male mice with no change in corneal shape, opacification of the cornea was still observed (n=22 eyes). Likewise, all opaque eyes showed neovascularization. Neovascularization of the cornea, regardless of morphological changes (n=28 eyes), was seen primarily in males, with <4% of females demonstrating neovascularization, likely associated with epithelial defects. A general bulging of the eye, without a cone-like change in curvature, was observed in both sexes (n=20). Evidence of corneal scarring, perforation, and hydrops (n=3) was only present in males. Localized corneal thinning was apparent in both male and female mice upon slit lamp exam, with 12 eyes showing centralized thinning and 21 eyes showing regional thinning in some other areas of the corneal surface.

**Figure 1.**
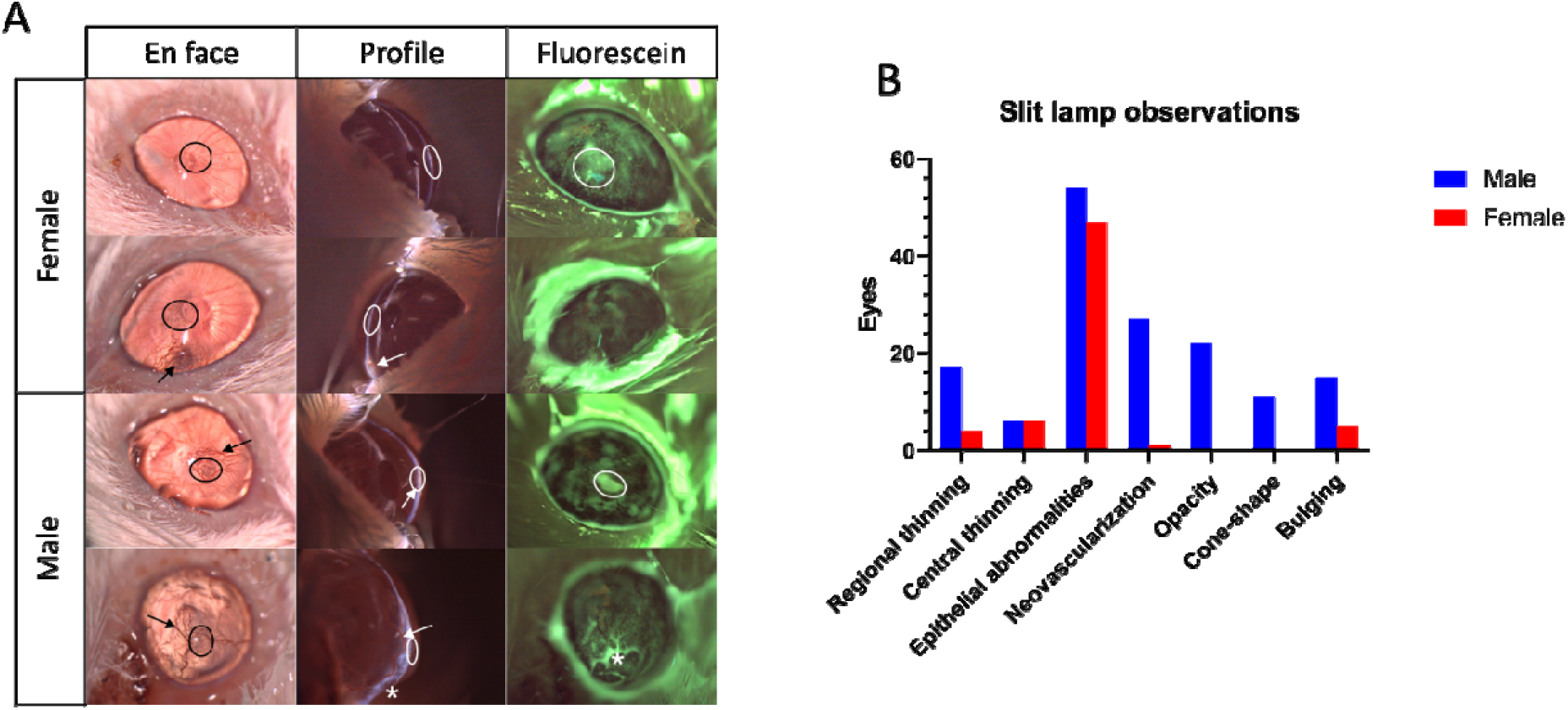
Representative slit lamp images of female and male SKC mouse corneas taken en face, profile, and en face with fluorescein (A). Circles indicate areas of epithelial abrasion or granulation, arrows indicate vascularization, and asterisks indicate suspected endothelial break or perforation. Summary of corneal phenotypes in male and female SKC mice as observed with slit lamp biomicroscopy (B).

### Corneal thickness

OCT-based pachymetry mapping of 8 mice showed changes to the corneal thickness in 14 of 16 eyes examined. Male SKC eyes (n=8) showed a wide variety of thickness changes, whereas all female eyes (n=8) examined showed some level of corneal thinning (Figure 2A). Half of female eyes (n=6) displayed a centralized thinning, like keratoconic corneas, and the other half (n=3 eyes) displayed a severe globalized thinning of the cornea, similar to keratoglobus. Males, on the other hand, displayed centralized thinning (n=3), centralized thickening (n=1), global thickening (n=4), and typical corneal thickness (n=2). One of the most severely affected eyes with a cone-like protrusion presented with a drastic central corneal thickening due to edema (Figure 2D). However, despite trends in pachymetry mapping, CCT measurements derived directly from SD-OCT scans did not reveal a statistically significant difference between males and females (t-test, p>0.05). The variance of CCT between groups, however, was statistically significant (p<0.0001) based on an F-test comparing standard deviations. This finding is consistent with the observation of large variability in thickness changes seen in males using pachymetry mapping.

**Figure 2.**
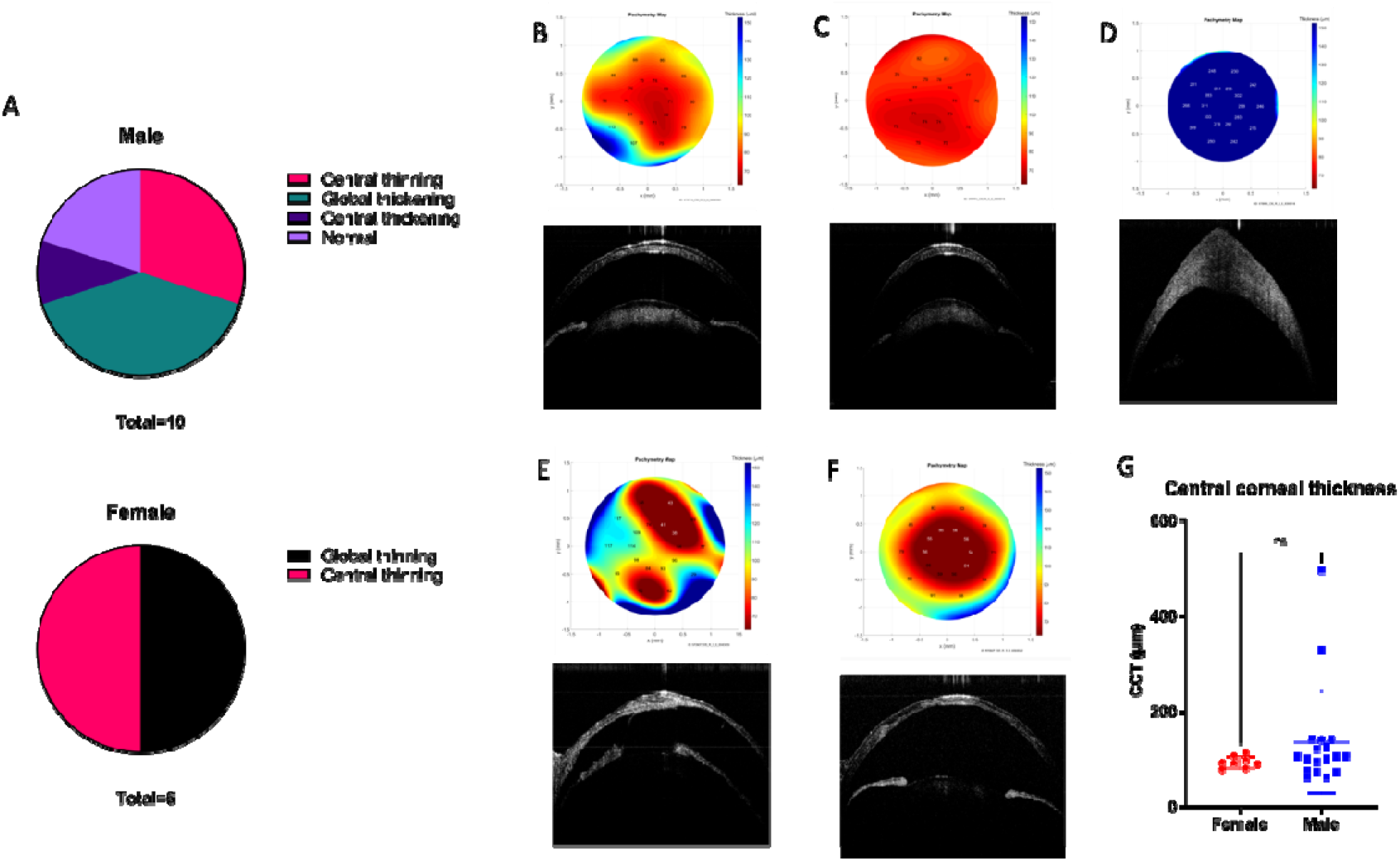
Summary of corneal thickness patterns observed via pachymetry mapping for male (n=10) and female (n=6) eyes (A). Pachymetry map with corresponding OCT cross-sectional images of female corneas (B, C), a cone-like male cornea (D), and non-cone-like male corneas (E, F). Comparison of central corneal thickness between female (n=8) and male (n=18) corneas with a student’s t-test (G).

### Histology and IVCM

Histological analysis of SKC corneas revealed several features such as epithelial abnormalities (n=11), breaks in the epithelial basement membrane (n=4), loosening of stromal compaction (n=15), increased cell density in the stroma (n=6), breaks in the endothelium (n=3), abnormal curvatures (n=10), and regional changes in corneal thickness (n=12). The most common histological finding observed in both female (n=7) and male (n=8) mouse eyes was stromal loosening, present in ∼68% of samples. Stromal loosening in SKC corneas exhibited variable severity, with a mild, anterior stromal loosening being most common (n=10) and a more severe manifestation representing ∼30% of stromal loosening in samples (n=5). Half of the samples assessed via H&E (n=11) demonstrated epithelial abnormalities, including thickening or thinning of the epithelium, irregular surface texture, abrasion, or breaks. Epithelial abnormalities were observed more frequently in males (n=7) than in females (n=4). Regional changes in corneal thickness were also common in over half of the samples examined (n=12). Regional changes in corneal thickness included central and localized corneal thinning or thickening.

In vivo cornea confocal microscopy of SKC corneas (n=9) revealed pathological features in both sexes, though males showed more severe characteristics. The epithelium presented with bright, reflective puncta in both sexes (n=5), though this was slightly more prevalent in male mice (n=3). All corneal stromas (n=9) demonstrated hyperreflectivity of the cytoplasm as well as enlarged and irregularly shaped nuclei in stromal keratocytes. Abnormal keratocyte nuclei included elongated, needle-like nuclei and spindle-like projections. Some of the male corneas (n=3) had a fiber-like appearance with few visible cell nuclei in the stroma. In male corneas only (n=3), blood vessels were clearly visible in the stroma with variable size and density. Abnormal appearance of stromal nerves was observed in both sexes, including enlargement, increased tortuosity, and non-uniform reflectivity of nerves (n=5). IV-CCM also revealed that basal epithelial and endothelial cell densities were significantly lower in males than in females.

### Immunofluorescence

Corneal sections from 4 mice (n=8 eyes) were stained with ZO-1, α-SMA, claudin-2, and SOD2. Staining of epithelial tight junctions with ZO-1 revealed an absent or weak expression of the protein in the basal epithelial layers in all corneas examined (n=8). Male corneas (n=4) demonstrated a weaker expression of ZO-1 overall when compared to most female corneas (n=3). One female cornea demonstrated a dramatic reduction in ZO-1 expression compared to all other female corneas (n=3), more similar to the presentation in male SKC mice. In contrast, claudin-2 expression in epithelial tight junctions was largely only evident in the basal strata of the epithelium (n=3), though some corneas demonstrated no claudin-2 staining at all (n=3). One female cornea and one male cornea demonstrated dramatic staining of claudin-2 in the entirety of the corneal epithelium, though fluorescence was strongest in the basal strata. SOD2 immunostaining was largely absent in the SKC corneas (n=7), though one male cornea with a more severe phenotype demonstrated staining throughout the corneal stroma. Finally, α-SMA staining was likewise rarely evident in SKC corneas, with half having no detectable expression (n=4), primarily the female corneas (n=3). Most male corneas (n=3) and one female cornea demonstrated α-SMA staining around the perimeters of elongated cells in the corneal stroma with variable levels of expression. In addition to stromal staining, one male cornea also exhibited α-SMA expression in Descemet’s membrane.

## Discussion

We sought to conduct an in-depth ophthalmic examination of the corneas of a mouse strain previously described as a spontaneously occurring, androgen-dependent keratoconus model [26]. We identified corneal phenotypes not previously reported in SKC mice through ophthalmic examination of 53 mice. In our assessment of these mice, we recapitulated several gross morphological characteristics originally described by Tachibana et al. (2002), including corneal bulging and steepened corneal angles in male mice [26]. We also detailed new characteristics of the corneal clarity, vascularization, epithelial integrity, and clinical signs of inflammation in both sexes that were not originally reported. Unlike original reports that abnormal corneal phenotypes were limited to male SKC mice, we observed gross abnormalities in female SKC mice. Major phenotypic findings were primarily related to epithelial integrity, corneal thickness, and corneal inflammation. Our study also suggested a lower penetrance of the cone-like phenotype (∼20%) than originally reported (∼33%)[26]. Overall, these findings suggest that unknown factors prevent severe phenotypic changes in female mice and contribute to the development of severe corneal changes in some, but not all, male SKC mice.

Epithelial defects and lesions were the most common phenotypic characteristics of SKC mice on slit lamp examination, observed in roughly 95% of the mice evaluated (Figure 1B). Male mice demonstrated a lower epithelial density on IV-CCM imaging than females (Figure 5A), potentially due to abrasion and epithelial cell apoptosis. In fact, IV-CCM images showed hyperreflective puncta in the epithelia of male and female mice (Figure 5C). These puncta may represent necrotic or apoptotic cells [29]. Immunofluorescence imaging revealed that epithelial tight junctions demonstrated major abnormalities in all male corneas and 2 female corneas (Figure 6). In these corneas, ZO-1 staining was either reduced or absent in the epithelium, usually with only the top-most 2-3 strata demonstrating any ZO-1 staining (Figure 6). ZO-1 serves an important role as a scaffold protein in epithelial tight junctions, binding other transmembrane and cytoplasmic components [30]. A decrease in ZO-1 expression in the corneal epithelium of SKC mice suggests that tight junction formation and assembly are impaired, likely increasing epithelial membrane permeability. Inversely, claudin-2 expression was elevated in half of the corneas imaged (n=3), especially in the middle and basal layers of the corneal epithelium (Figure 6). Claudin-2 is a component of epithelial tight junctions, specifically those of “leaky” epithelia [31, 32]. Claudin-2 forms cation-selective, water-permeable paracellular channels in tight junctions, mediating water transport across the epithelium [31]. The elevated staining of claudin-2 in SKC corneas suggests a loss of typical tight junction structure and barrier function, creating a “leaky” corneal epithelium.

A range of changes to corneal thickness were observed, including localized thinning or thickening (Figure 2A). Notably, there were cases of corneal thickening, which corresponded to some of the cone-like and bulging corneas (Figure 2D). This thickening was likely caused by corneal edema because of corneal perforation or other endothelial defects, which is inconsistent with human KC. Histological analysis revealed stromal loosening (Figure 3C, D) and endothelial adhesion to the iris (Figure 3F), indicative of corneal edema and perforation. Reduced endothelial cell density in male SKC mice, based on IV-CCM (Figure 5A), may account for the increase in corneal thickness due to stromal edema, though the impetus behind the reduction in cell density is not clear. Pachymetry mapping also revealed universal thinning of some female corneas (Figure 2A, C), similar to keratoglobus, the cause of which is not clear but appears to be endogenous. However, despite visual trends observed in pachymetry mapping, CCT measurements derived from OCT scans showed no statistical significance between males and females (Figure 2G). Though the CCT measurements do not factor in areas of non-centralized thinning or the central corneal thickness relative to the peripheral thickness of each cornea, both of which are better represented by pachymetry mapping. Additionally, variations in corneal thickness patterns do not allow for an accurate comparison with human KC.

**Figure 3.**
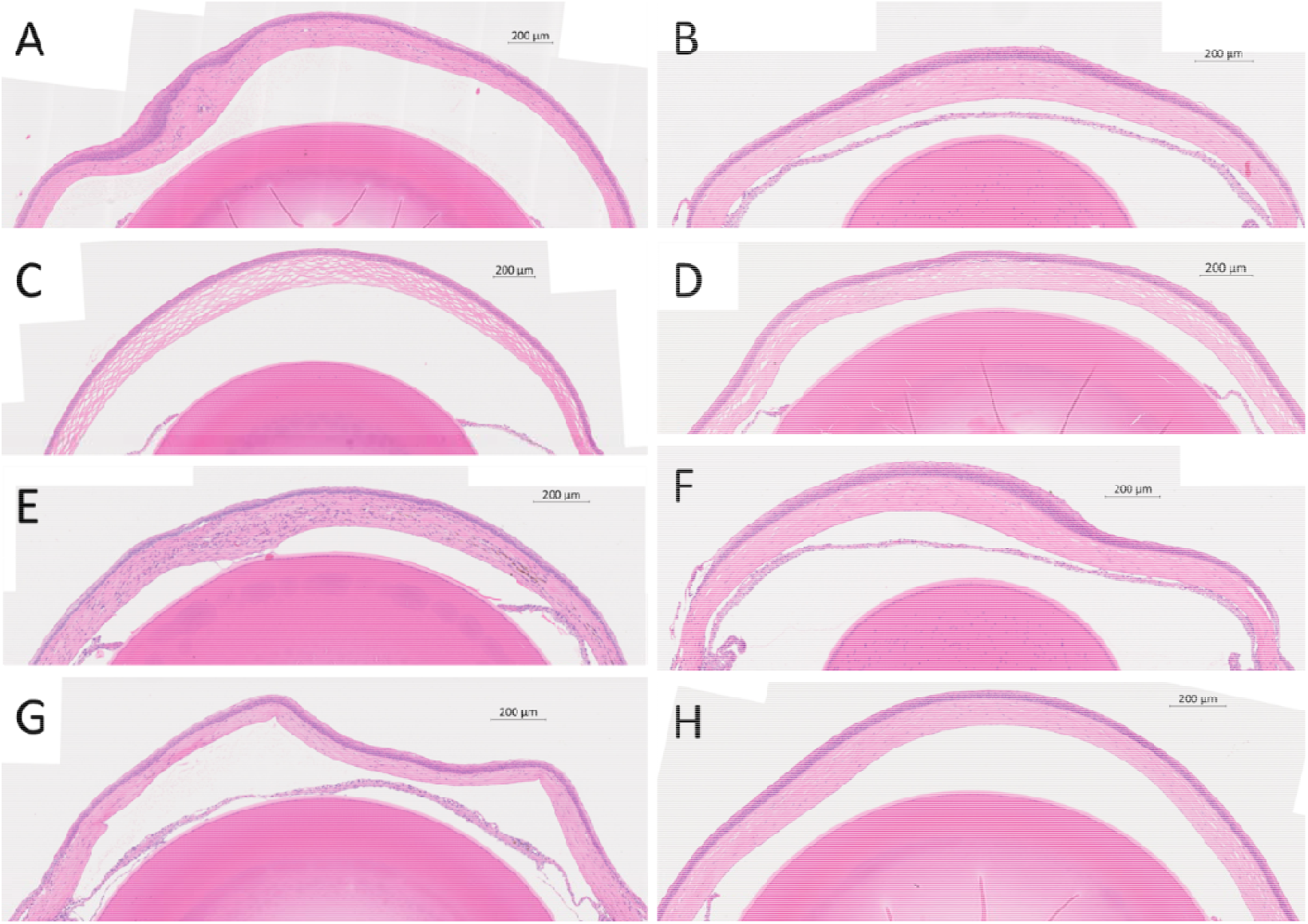
H&E images of male (A, C, E, G) and female (B, D, F, H) SKC mouse eyes. Scale bar: 200µm.

**Figure 4.**
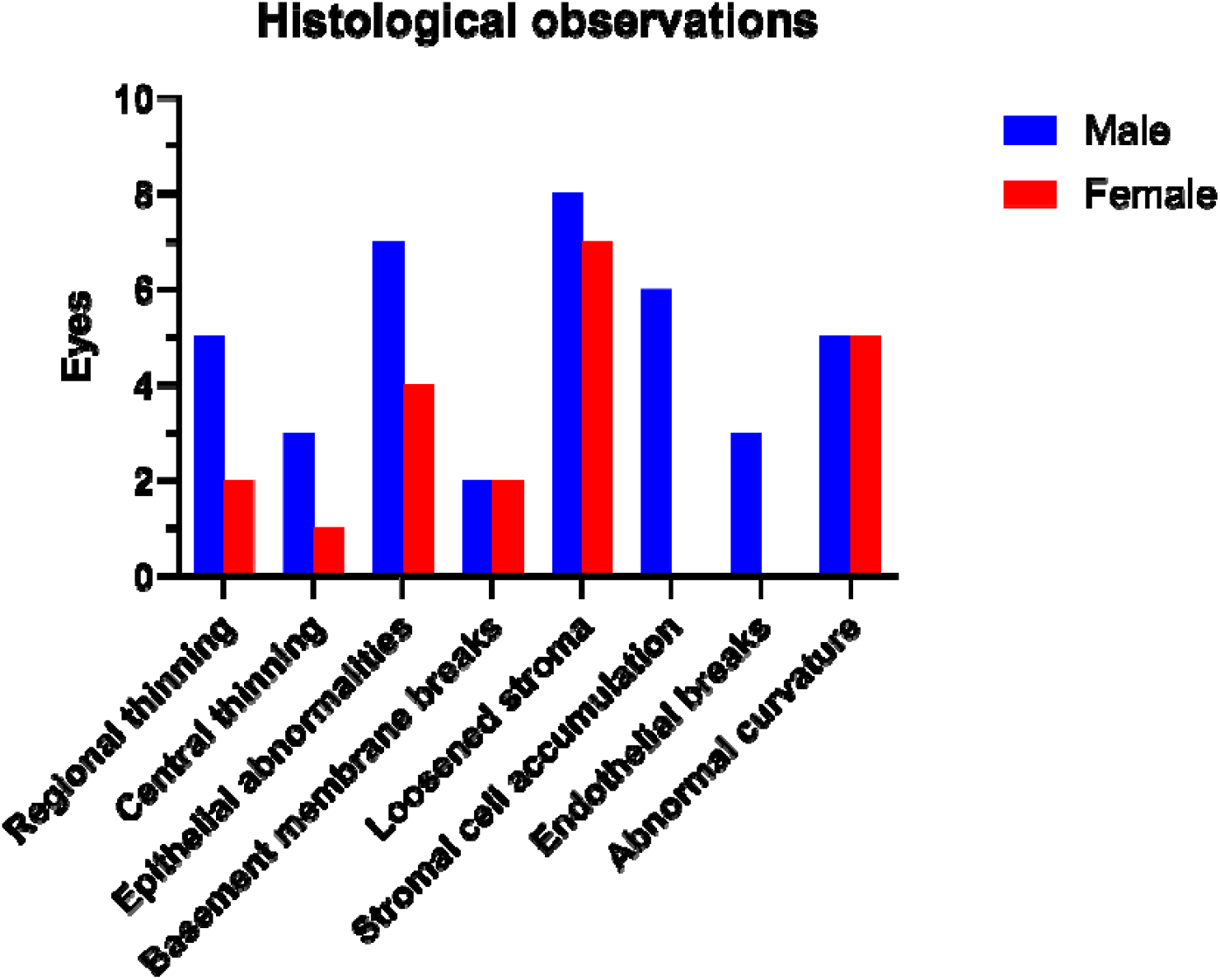
Summary of histological abnormalities observed in male and female SKC mice.

Additional clinical indicators of corneal inflammation and fibrosis were apparent upon slit lamp examination, including neovascularization, scarring, and hydrops (Figure 1A). These characteristics were primarily observed in males, though some females also exhibited neovascularization (Figure 1B). Blood vessels were visible in male corneas via IV-CCM (Figure 5F), which showed vascularization of the stroma with size and density of blood vessels largely reflective of gross corneal phenotypes. Three male corneas and one female cornea displayed α-SMA expression via immunofluorescence (Figure 6). Expression of α-SMA indicates that keratocytes in the stroma are undergoing myofibroblast transformation, which primarily occurs in response to corneal injury and infection [33]. Myofibroblasts are major contributors to stromal scarring and fibrosis under dysregulated conditions, during which they develop a contractile phenotype and produce disordered ECM [33]. Some studies have shown that α-SMA expression is increased in KC corneal cells and tissues [34, 35]. This finding recapitulates observations of abnormal keratocyte morphology and ECM organization seen in IV-CCM, including a fiber-like appearance of keratocyte cytoplasm in the male corneal stroma, elongated or needle-like nuclei, and spindle-like projections (Figure 5C, E). Previously, Quantock et al. (2003) stated that the cone-like corneal phenotype of SKC mice may be secondary to keratitis [27]. Many of our findings substantiate this in mice with more severely affected corneas, as evidenced by severe corneal edema, neovascularization, and epithelial damage in gross observations, histology, immunofluorescence, and confocal microscopy analysis. Given that human keratoconus is not characterized by clinical signs of ocular inflammation such as swelling and redness, the SKC corneal phenotype is inconsistent with KC [36]. Though molecular inflammatory factors may play a role in KC pathogenesis [37].

**Figure 5.**
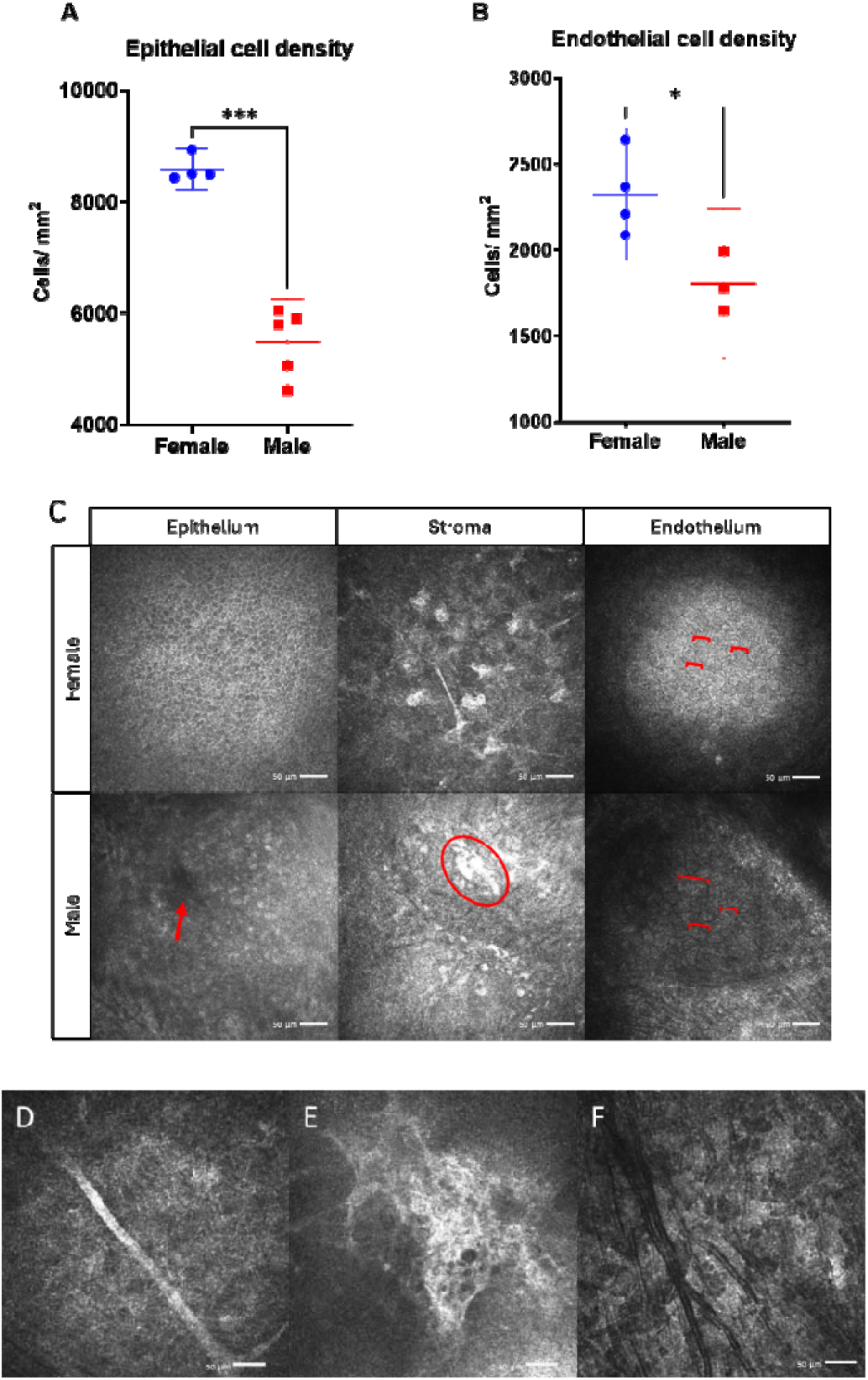
Comparison of epithelial (A) and endothelial (B) cell density calculations in female and male SKC mice based on IV-CCM images. Representative IV-CCM images of the corneal epithelial, stromal, and endothelial layers in female and male mice (C); arrows depict the region of epithelial cell gapping, circles identify hyperreactive stromal cell nuclei, and brackets depict the circumference of endothelial cells. Appearance of an abnormal stromal nerve in a female cornea (D). Presence of fibrotic stromal keratocytes in a male cornea (E). Vascularization is evident in the stroma of a male cornea (F). Scale bar: 50µm.

**Figure 6.**
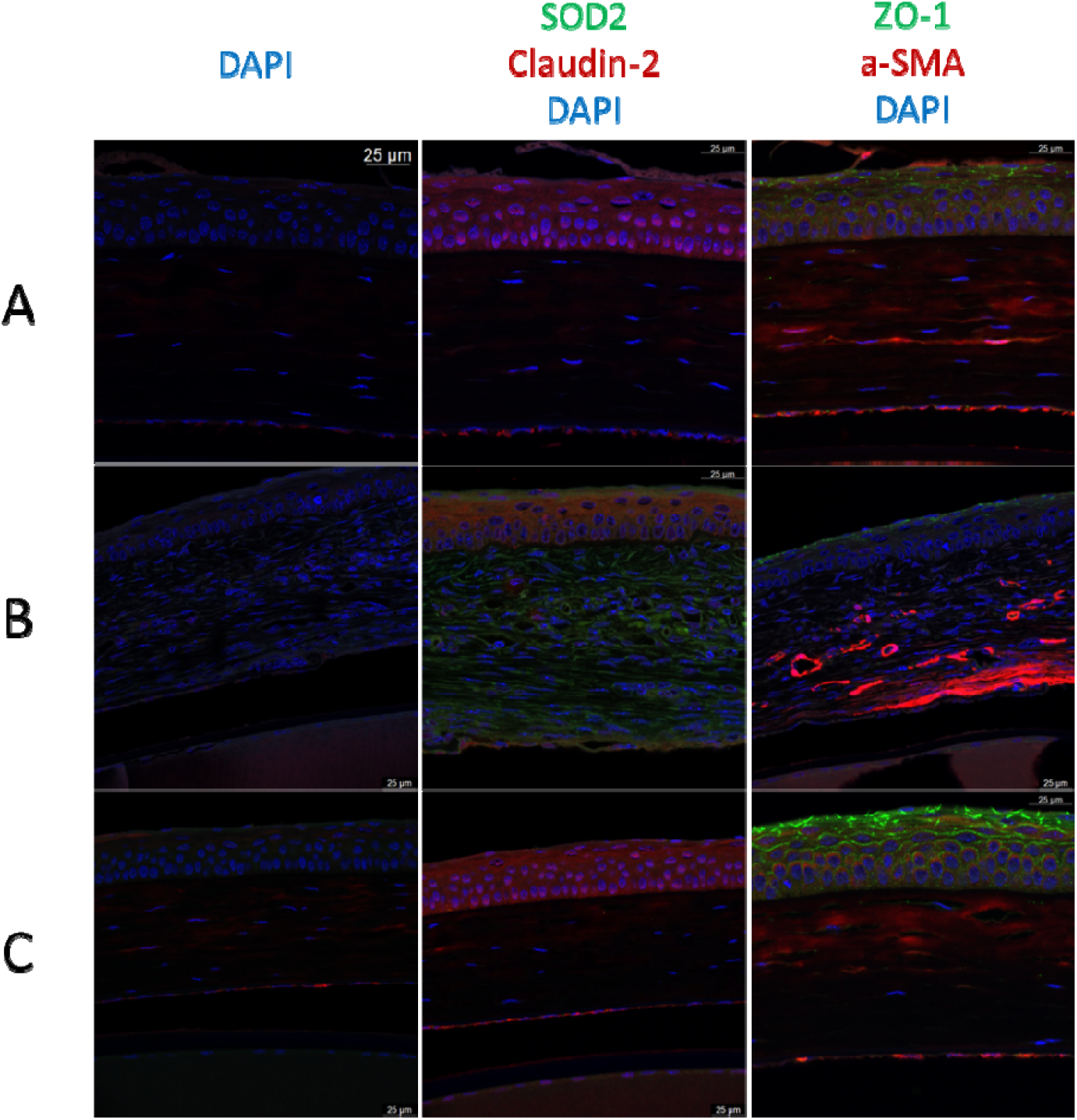
Immunofluorescence images of separate male corneas (A, B) and female cornea (C) at 40x magnification. Column 1 depicts no-primary antibody negative controls for each corresponding sample. Column 2 depicts staining against superoxide dismutase 2 (SOD2, green) and claudin 2 (red) proteins. Column 3 depicts staining against zona-occludens 1 (ZO-1, green) and alpha-smooth muscle actin (α-SMA, red). Nuclei are stained with DAPI (blue) in all images. Scale bar: 25µm.

In addition to the corneal phenotypes, we also observed evidence of dermatitis and scratching in the mice. Mice with more severe corneal abnormalities exhibited more severe dermatitis and self-scratching behavior, including around the eyes. It is likely that this more aggravated scratching behavior exacerbates corneal inflammation. Atopic dermatitis has previously been associated with keratoconus, with a more recent study asserting a causal relationship between the two [36, 38-40]. Previously, corneal abnormalities were observed in a mouse model of atopic dermatitis: the NC/Nga mouse [41]. The gross appearance of the skin of NC/Nga mice closely resembles that of SKC mice, and they also exhibited eye-scratching behaviors [41]. Corneal abnormalities in the NC/Nga included epithelial thinning, abnormal curvature, subepithelial deposition, and stromal vascularization similar to some SKC mouse characteristics [41]. Although we have not validated that the dermatitis observed in SKC mice is atopic dermatitis rather than other dermatological conditions, it is possible that SKC mouse could reveal factors that contribute to the association between KC and atopic dermatitis, which requires further investigation.

The enduring lack of an in vivo experimental model to study keratoconus continues to limit progress in the field. Although continued efforts are underway to generate a genetic animal model for KC, the multigenic nature of KC makes a spontaneously occurring phenotype ideal. However, our assessment of the SKC mouse suggests that several characteristics are not only inconsistent with KC, but also indicative of pathological processes beyond KC. One important limitation of this study is the lack of a non-SKC control strain included in our analysis. The different phenotypes we identified could also be impacted by the different environment in Augusta and the animal facility compared to those in Japan many years ago.

It may be beneficial for future follow-up studies to investigate the extent of molecular changes in female and mildly affected male SKC corneas compared with non-SKC mice, to assess whether these cases may be representative of human keratoconus at mild or subclinical stages, and to evaluate the molecular mechanisms at play.

## References

1. Rabinowitz, Y.S., Keratoconus. Surv Ophthalmol, 1998. 42(4): p. 297–319.

2. Santodomingo-Rubido, J., et al., Keratoconus: An updated review. Contact Lens and Anterior Eye, 2022. 45(3): p. 101559.

3. Espandar, L. and J. Meyer, Keratoconus: overview and update on treatment. Middle East Afr J Ophthalmol, 2010. 17(1): p. 15–20.

4. Torres Netto, E.A., et al., Prevalence of keratoconus in paediatric patients in Riyadh, Saudi Arabia. Br J Ophthalmol, 2018. 102(10): p. 1436–1441.

5. Gorskova, E.N. and E.N. Sevost’ianov, [Epidemiology of keratoconus in the Urals]. Vestnik oftalmologii, 1998. 114(4): p. 38–40.

6. Gomes, J.A.P., P.F. Rodrigues, and L.L. Lamazales, Keratoconus epidemiology: A review. Saudi J Ophthalmol, 2022. 36(1): p. 3–6.

7. Singh, R.B., U.P.S. Parmar, and V. Jhanji, Prevalence and Economic Burden of Keratoconus in the United States. American Journal of Ophthalmology, 2024. 259: p. 71–78.

8. Parker, J.S., K. van Dijk, and G.R. Melles, Treatment options for advanced keratoconus: A review. Surv Ophthalmol, 2015. 60(5): p. 459–80.

9. Wollensak, G., E. Spoerl, and T. Seiler, Riboflavin/ultraviolet-a-induced collagen crosslinking for the treatment of keratoconus. Am J Ophthalmol, 2003. 135(5): p. 620–7.

10. Ashwin, P.T. and P.J. McDonnell, Collagen cross-linkage: a comprehensive review and directions for future research. British Journal of Ophthalmology, 2010. 94(8): p. 965–970.

11. Galvis, V., et al., Patient selection for corneal collagen cross-linking: an updated review. Clin Ophthalmol, 2017. 11: p. 657–668.

12. Wang, Y., et al., Genetic epidemiological study of keratoconus: evidence for major gene determination. Am J Med Genet, 2000. 93(5): p. 403–9.

13. Jaskiewicz, K., et al., Non-allergic eye rubbing is a major behavioral risk factor for keratoconus. PLoS One, 2023. 18(4): p. e0284454.

14. Hashemi, H., et al., The Prevalence and Risk Factors for Keratoconus: A Systematic Review and Meta-Analysis. Cornea, 2020. 39(2): p. 263–270.

15. Fink, B.A., et al., Differences in keratoconus as a function of gender. Am J Ophthalmol, 2005. 140(3): p. 459–68.

16. Delic, N.C., et al., Damaging Effects of Ultraviolet Radiation on the Cornea. Photochem Photobiol, 2017. 93(4): p. 920–929.

17. Bykhovskaya, Y. and Y.S. Rabinowitz, Update on the genetics of keratoconus. Exp Eye Res, 2021. 202: p. 108398.

18. Anitha, V., et al., Pediatric keratoconus - Current perspectives and clinical challenges. Indian J Ophthalmol, 2021. 69(2): p. 214–225.

19. DelMonte, D.W. and T. Kim, Anatomy and physiology of the cornea. Journal of Cataract & Refractive Surgery, 2011. 37(3): p. 588–598.

20. Giugliani, R., et al., Mucopolysaccharidosis I, II, and VI: Brief review and guidelines for treatment. Genet Mol Biol, 2010. 33(4): p. 589–604.

21. Boote, C., et al., Collagen fibrils appear more closely packed in the prepupillary cornea: optical and biomechanical implications. Invest Ophthalmol Vis Sci, 2003. 44(7): p. 2941– 8.

22. Hovakimyan, M., et al., Morphological analysis of quiescent and activated keratocytes: a review of ex vivo and in vivo findings. Curr Eye Res, 2014. 39(12): p. 1129–44.

23. Khaled, M.L., et al., Molecular and Histopathological Changes Associated with Keratoconus. BioMed Research International, 2017. 2017: p. 7803029.

24. Hadvina, R., A. Estes, and Y. Liu, Animal Models for the Study of Keratoconus. Cells, 2023. 12(23): p. 2681.

25. Loiseau, A., et al., Animal Models in Eye Research: Focus on Corneal Pathologies. International Journal of Molecular Sciences, 2023. 24(23): p. 16661.

26. Tachibana, M., et al., Androgen-Dependent Hereditary Mouse Keratoconus: Linkage to an MHC Region. Investigative Ophthalmology & Visual Science, 2002. 43(1): p. 51–57.

27. Quantock, A.J., et al., Annulus of collagen fibrils in mouse cornea and structural matrix alterations in a murine-specific keratopathy. Invest Ophthalmol Vis Sci, 2003. 44(5): p. 1906–11.

28. Liu, A.S., et al., Topography and pachymetry maps for mouse corneas using optical coherence tomography. Exp Eye Res, 2020. 190: p. 107868.

29. Jalbert, I., et al., In vivo confocal microscopy of the human cornea. Br J Ophthalmol, 2003. 87(2): p. 225–36.

30. Van Itallie, C.M., et al., ZO-1 Stabilizes the Tight Junction Solute Barrier through Coupling to the Perijunctional Cytoskeleton. Molecular Biology of the Cell, 2009. 20(17): p. 3930–3940.

31. Rosenthal, R., et al., Claudin-2, a component of the tight junction, forms a paracellular water channel. Journal of Cell Science, 2010. 123(11): p. 1913–1921.

32. Venugopal, S., S. Anwer, and K. Szászi, Claudin-2: Roles beyond Permeability Functions. International Journal of Molecular Sciences, 2019. 20(22): p. 5655.

33. Wilson, S.E., Corneal myofibroblasts and fibrosis. Experimental Eye Research, 2020. 201: p. 108272.

34. Karamichos, D., et al., Novel in Vitro Model for Keratoconus Disease. J Funct Biomater, 2012. 3(4): p. 760–775.

35. Wang, Y.N., et al., Expression of visual system homeobox 1 in human keratoconus. Int J Ophthalmol, 2019. 12(2): p. 201–206.

36. Rabinowitz, Y.S., Keratoconus. Survey of Ophthalmology, 1998. 42(4): p. 297–319.

37. Galvis, V., et al., Keratoconus: an inflammatory disorder? Eye, 2015. 29(7): p. 843–859.

38. Pietruszyńska, M., et al., Ophthalmic manifestations of atopic dermatitis. Postepy Dermatol Alergol, 2020. 37(2): p. 174–179.

39. Harrison, R.J., et al., Association between keratoconus and atopy. Br J Ophthalmol, 1989. 73(10): p. 816–22.

40. Chang, Y., et al., Causal Association Between Atopic Dermatitis and Keratoconus: A Mendelian Randomization Study. Transl Vis Sci Technol, 2024. 13(9): p. 13.

41. Ebihara, N., et al., Corneal abnormalities in the NC/Nga mouse: an atopic dermatitis model. Cornea, 2008. 27(8): p. 923–9.

